# MOLECULAR CORRELATE OF MOUSE EXECUTIVE FUNCTION. TOP-DOWN AND BOTTOM-UP COMPLEMENTATIONS BY PRESYNAPTIC VERTEBRATE BRAIN-SPECIFIC Ntng GENE PARALOGS

**DOI:** 10.1101/139444

**Authors:** Pavel Prosselkov, Qi Zhang, Hiromichi Goto, Denis Polygalov, Thomas J. McHugh, Shigeyoshi Itohara

## Abstract

Executive function (EF) is a regulatory construct of learning and general cognitive abilities. Genetic variations underlying the architecture of cognitive phenotypes are likely to affect EF and associated behaviors. Mice lacking one of *Ntng* gene paralogs, encoding the vertebrate brain-specific presynaptic Netrin-G proteins, exhibit prominent deficits in the EF control. Brain areas responsible for gating the bottom-up and top-down information flows differentially express *Ntng1* and *Ntng2*, distinguishing neuronal circuits involved in perception and cognition. As a result, high and low cognitive demand tasks (HCD and LCD, respectively) modulate *Ntng1* and *Ntng2* associations either with attention and impulsivity (AI) or working memory (WM), in a complementary manner. During the LCD *Ntng2*supported neuronal gating of AI and WM dominates over the *Ntng1*-associated circuits. This is reversed during the HCD, when the EF requires a larger contribution of cognitive control, supported by *Ntng1*, over the *Ntng2* pathways. Since human *NTNG* orthologs have been reported to affect human IQ (1), and an array of neurological disorders (2), we believe that mouse *Ntng* gene paralogs serve an analogous role but influencing brain executive functioning.

## INTRODUCTION

Executive function (EF) is a heterogeneous construct that can be viewed as a set of processes executively supervising cognitive behaviors (3). EF is an umbrella term for working memory (WM), attention and impulsivity (AI), and response inhibition, and is thought to account for the variance in cognitive performance (4). WM, due to its storage and processing components, is viewed as a bimodal flexible system of a limited capacity. Since WM maintains current information and simultaneously supports its execution, as a latent factor underlying intelligence (5), it has been termed as “the central executive” (6) attention-controlling system dependent on consciousness (7). However an awareness-independent model has been also proposed (8,9). General learning (Ln) ability depends on attention and WM interaction (10) as well as perception, the causal and informational ground for the higher cognitive functions (11). Perception guides our thinking about and acting upon the world and serves as an input to cognition, via a short-term memory mediated interactions (12). A possible mechanism linking perception and cognition would be attention (13).

Perception (bottom-up) and cognition (top-down) have been historically viewed as independently operating encapsulating domains. Such embodiment has paved a ground for the view that perceptual experiences can be influenced by cognitive state (for references see 14), consequently elaborated into the brain predictive coding approach currently dominating cognitive neuroscience (15), and positing that attention is a property of brain computation network (16). However this has been challenged by the opposite opinion that “cognition does not affect perception” (17). Regardless whether or not such a cognitive-sensory dichotomy exists, herein we view perception and cognition as two main information streams the EF exerts its actions upon, possibly through active association.

We have previously described the function of two vertebrate-specific brain-expressed presynaptic gene paralogs, *NTNG1* and *NTNG2*, complementary affecting verbal comprehension and WM in human subjects, that underwent an accelerated evolution in primates and extinct hominins (1). This pair of genes has been also implicated in the phenomena of antagonistic pleiotropy, a trade-off between the evolution-driven cognitive function elaboration and an array of concomitant neuropathologies, rendering the human brain phenotypically fragile (2). *Ntngs* also complementary diversify the mouse behavior (18).

Despite the fact that EF abrogation is a major determinant of problem behavior and disability in neuropsychiatric disorders (19), the genetics underlying EF remains elusive with no causative vector agents (e.g. genes) have yet been reported. Herein we show that *NTNG* paralogs affecting human IQ also affect mouse learning and brain executive functioning.

## RESULTS

### Randomizing mouse genotypes in a search for genotype-phenotype interactions

We used a non-parametric data analysis approach for two behavioral paradigms: 5-choice serial reaction time task (5-CSRTT, 20), and radial arm maze (RAM), to measure selective attention and impulsivity (AI), and spatial working memory (WM), respectively, in *Ntng1*^-/-^and *Ntng2*^-/-^mice. We calculated mouse genotype-independent ranking (as for a mixed population), and the rank variance (as a proportion of variance explained, PVE) for each behavioral parameter and a global rank for each paradigm. This allowed us to avoid common in a behavior-reporting literature a genotype-attributed single parameter reporting bias via the data “cherry-picking” satisfying common standards of statistical significance tests (e.g. 18, see Supplementary Figures 1 (SF1) and 2 (SF2) for the same data being processed in a traditional way). This also permitted us to compare the observed phenotypes between both paradigms for the genetically independent groups of mice, simultaneously searching for potential interactions among them. We were able to follow the dynamics of the behavioral heterogeneity and to deduce inferences between the mouse phenotypic and genotypic traits interaction affecting executive function (EF).

### Affected AI for both *Ntng* paralogs, and WM for the *Ntng2* gene, modulated by the cognitive demand

5-CSRTT data (ST1-1) show that *Ntng1*^-/-^population of mice is characterised by a large span of the ranks varaince (PVE>90%) occupying not only bottom 4 but also top 4 rank positions and outcompeting their wild type littermates (Fig.1(A-D)-1). *Ntng1* ablation generates mice with both strong proficit and deficit of AI, extending beyond a single affected parameter estimate (Fig.1C,G), but with the averaged rank per a genotype undistinguishable of that of their wild type littermates, and more than 90% of the variance attributable to *Ntng1*^-/^genotype (Fig.1A-1). A higher cognitive demand task phase (HCD) reduces the rank variance down to 76% but at the expense of a lower rank (Fig.1E-1). During the low cognitive demand task phase (LCD), opposite to *Ntng1*^-/-^mice, *Ntng2*^-/-^subjects’ rank is twice lower comparing to their genetically unmodified littermates with the rank variances almost identical (Fig.1A-2) but later changed during the HCD (Fig.1E-2).

**Figure 1.**
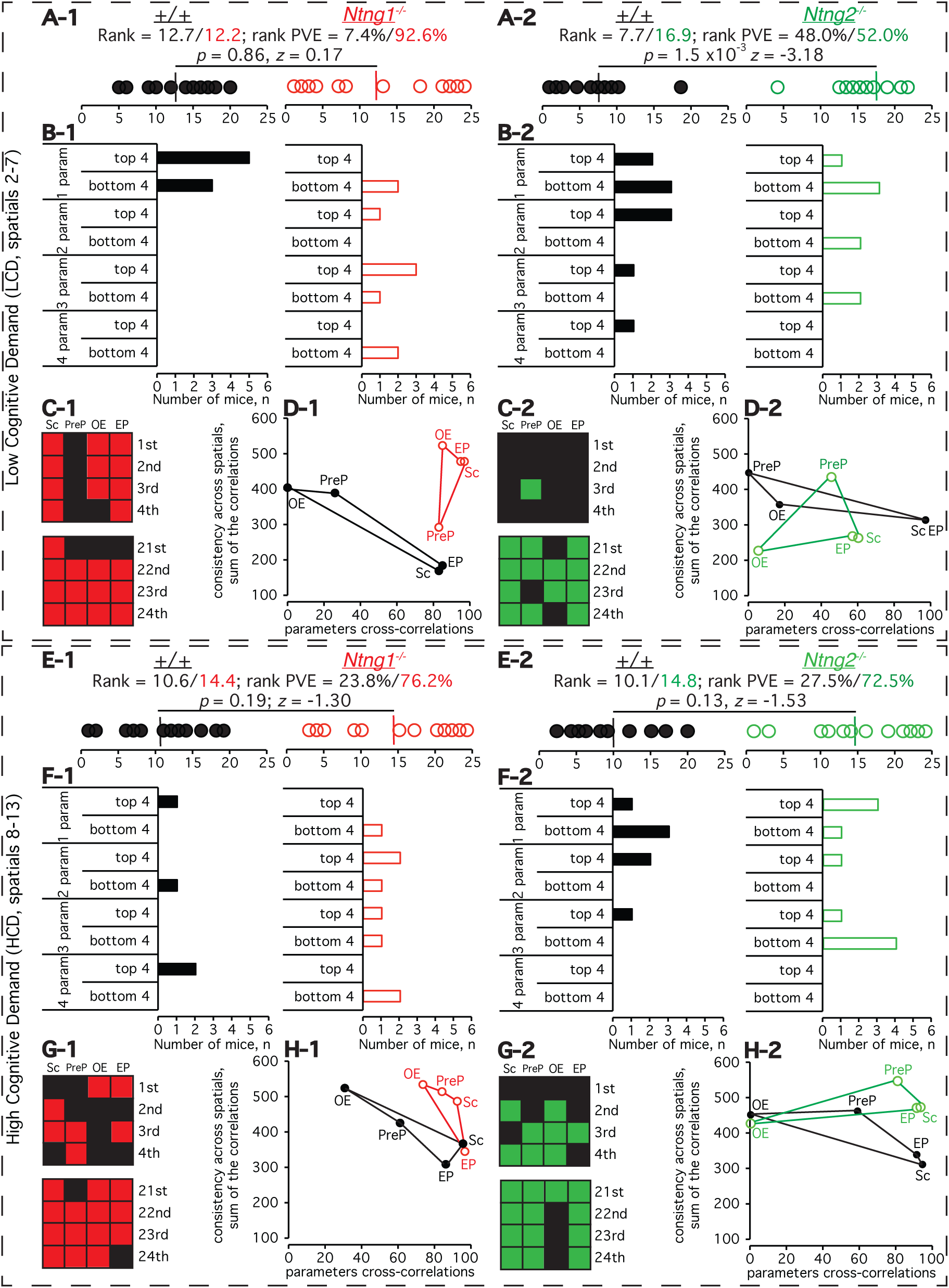
Attention and Impulsivity (AI) estimate and the effect of cognitive demand by the analysis of rank and its variance for *Ntng1*^-/-^and *Ntng2*^-/-^mice. A,E. Mouse ranks and rank PVE (proportion of variance explained) based on four parameter rank measures (SF1) as detailed in ST1-1 (for *Ntng1*^-/-^) and ST1-2 (for *Ntng2*^-/-^). The rank sorting was done in a genotype-independent manner treating all micetogether. Ranking for each out of four parameters was done independently of other parameters with afinal re-ranking of the ranks sum to generate the final rank (shown). In case of an equal sum of the ranks, the mice were given identical ranks. PVE was calculated as a square of within genotype rank variance divided on the sum of each genotype variances squares multiplied on 100%. **B,F**. Mouse rank distribution across one-to-four parameters as top 4 and bottom 4 performers. **C,G**. Genotype-specific placing among the mice. **D,H**. Behavioral consistency of mice across the sessions (*y* axis, sum of r^2^ correlations of a single session ranks vs. final ranks for each mouse across the sessions) and behavioral parameter cross-correlations (*x* axis, the r^2^ correlation of a parameter final ranking *vs*. final ranking for all 4 parameters). The gene ablation-specific phenotype severity can be assessed visually by matching each parameter-corresponding verteces of the obtained quadruples. *p* value represents a Wilcoxon rank sum test.

Robustness of the WM deficit upon *Ntng2* depletion in mice is the most prominently evidenced by the bottom 4 mouse ranks with 12/13 out of 16 being occupied by the knockout mice (Fig.2C-2,G-2) and by low behavioral consistency across the sessions and parameters cross-correlations (Fig.2H-2) during the HCD. At the same time, the absence of *Ntng1* in mice affected only the LCD sessions performance (Fig.2D-1) but did not render them behaviorally distinguishable from the wild type littermates during the HCD (Fig.2H-1). s

**Figure 2.**
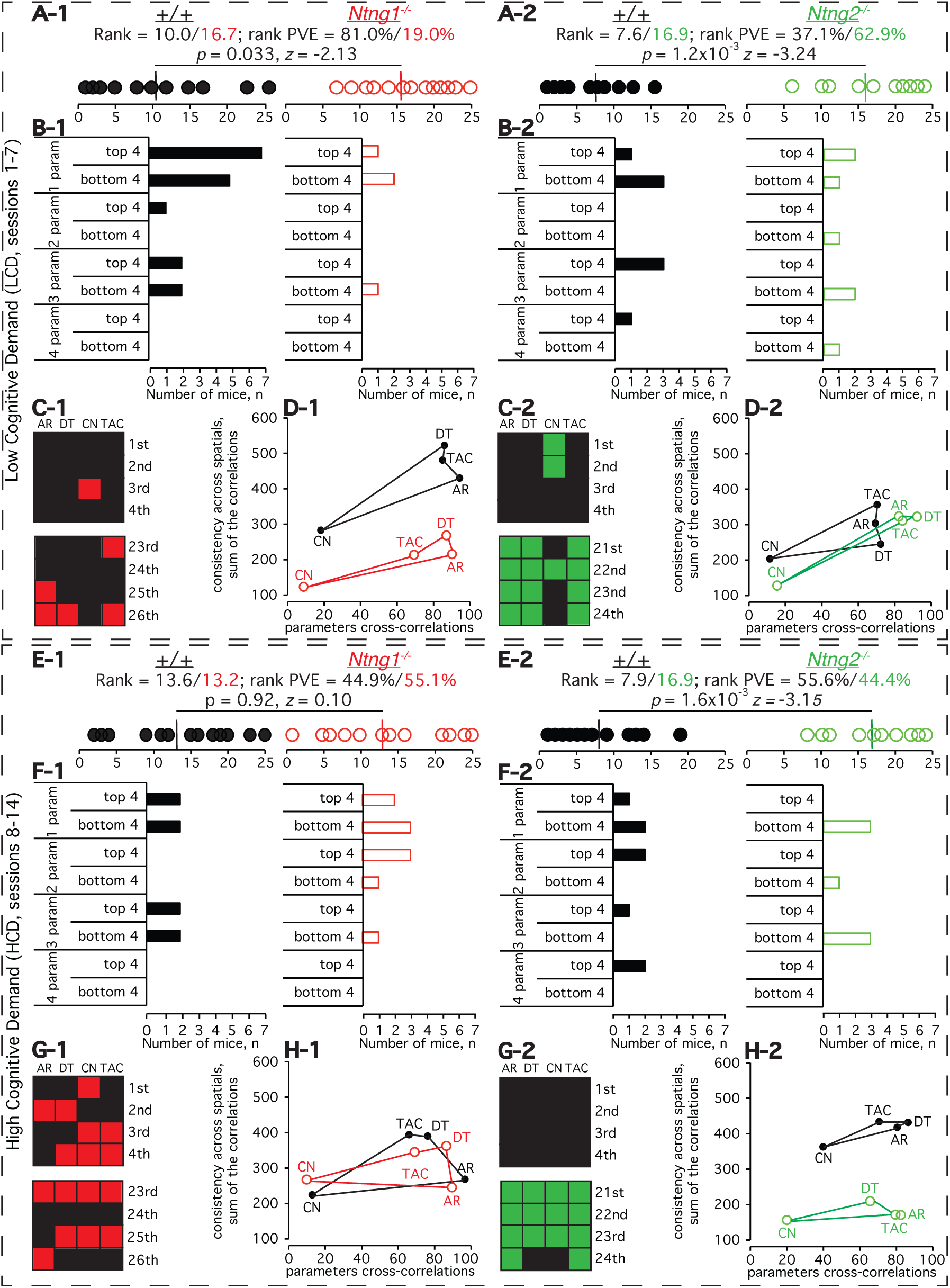
Working memory (WM) estimate and the effect of cognitive demand by the analysis of rank and its variance for *Ntng1*^-/-^and *Ntng2*^-/-^mice. **A,E**. Mouse ranks and rank PVE (proportion of variance explained) based on four parameter rank measures (SF2) as detailed in ST2-1 (for *Ntng1*^-/-^) and ST2-2 (for *Ntng2*^-/-^). The rank sorting was done in a genotype-independent manner treating all mice together. Ranking for each out of four parameters was done independently of other parameters with a final re-ranking of the ranks sum to generate the final rank (shown). In case of an equal sum of the ranks, the mice were given identical ranks. PVE was calculated as a square of within genotype rank variance divided on the sum of each genotype variances squares multiplied on 100%. **B,F**. Mice rank distribution across one-to-four parameters as top 4 and bottom 4 performers. **C,G**. Genotype-specific placing among the mice. **D,H**. Behavioral consistency of mice across the sessions (*y* axis, sum of r^2^ correlations of a single session ranks *vs*. final ranks for each mouse across the sessions) and behavioral parameter cross-correlations (*x* axis, the r^2^ correlation of a parameter final ranking *vs*. final ranking for all 4 parameters). The gene ablation-specific phenotype severity can be assessed visually by matching each parameter-corresponding vertexes of the obtained quadruples. *p* value represents a Wilcoxon rank sum test.

### Proficit and deficit in learning associated with the *Ntng*^-/-^genotypes

The complementary segregation of *Ntng*^-/-^gene paralogs-associated behavioral phenotypes within the distinct modules of EF (Fig.3; ST1 and ST2) has prompted us to analyse the operant conditioning learning (ST1-1 and ST1-2 (Sc_spatial1, Ln)) by mice, assuming that AI and WM may interact. And indeed, *Ntng1*^-/-^ mice outperform their genetically unmodified littermates learning faster during the LCD (Fig.4(A,B)-1, LCD) but are unable to sustainably cope with the growing cognitive demand (Fig.4(A,B)-1, HCD). At the same time, *Ntng2*^-/-^mice display a prominent deficit of Ln (Fig.4(A,B)-2,LCD), which is becoming stronger with the growing demand to succeed (Fig.4(A,B)-2, HCD). In overall, the pattern of Ln behavior caused by the genetic ablation of either of *Ntngs* completely matches that of WM testing on the RAM (Fig.2), summarised in Fig.3. The contribution of AI to the Ln deficit is demonstrated by the rank correlations of Ln *vs*. AI (from Fig.1) which is stronger during the LCD for both *Ntng* genetically ablated mouse populations (Fig.4C-1,2).

**Figure 3.**
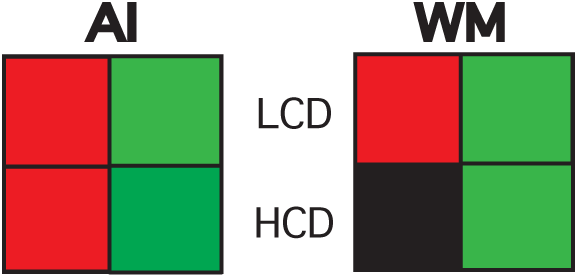
Summary of the EF behavioral phenotypes associated with either *Ntng1*^-/-^or *Ntng2*^-/-^gene paralogs ablation. See Fig.1 and Fig.2 for details.

**Figure 4.**
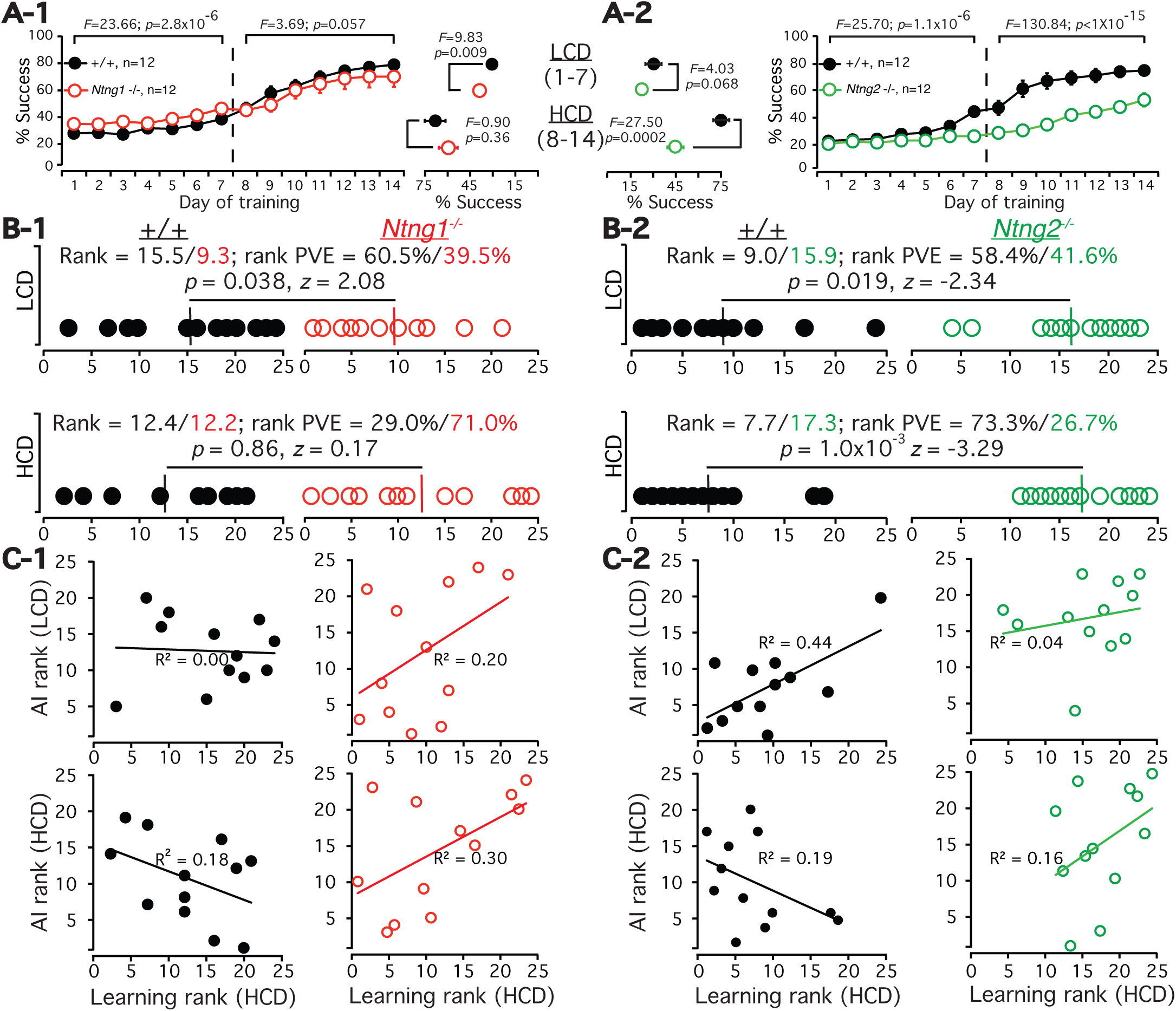
Operant conditioning (5-CSRTT) Learning (Ln) by *Ntng1*^-/-^and *Ntng2*^-/-^mice. **A**. Learning curves for the operant conditioning learning (reward collection) over the training period (spatial 1 of 5-CSRTT) with averaged performance behavior for the days (1-7) and (8-14), middle panel, defined as low cognitive demand (LCD) and high cognitive demand (HCD) sessions, respectively. One and two-way ANOVA was used for the statistics. **B**. Ranks and PVE comparisons over the LCD and HCD. The rank sorting was done in a genotype-independent manner, similar to Fig.1 and Fig.2, but using only one parameter, success (Sc), see ST2-1 and ST2-2 (Ln). Rank statistics was by Wilcoxon rank sum test. **C**. Learning (Ln) *vs*. attention and impulsivity (AI) rank correlations (from Fig.1A-1,2).

### Complementary expression of *Ntng* paralogs in the brain and their interaction

The robust phenotype of the abrogated EF for both *Ntng* gene paralogs affecting either AI or WM, or both, is supported by the predominant expression of both genes within the information processing brain areas, complementary sequestering them either within bottom-up (for *Ntng1*) and top-down (for *Ntng2*) neuronal pathways (Fig.5A-C). The presented hierarchy for the *Ntng* paralogs brain distribution is supported by two times lower level of the *Ntng2* expression under *Ntng1*^-/-^background (Fig.5D-2), with no effect on *Ntng1* expression when *Ntng2* is absent (Fig.5D-1) in the life-long cognitively trained in senile mice (randomly selected from ST-1 and ST-2).

**Figure 5.**
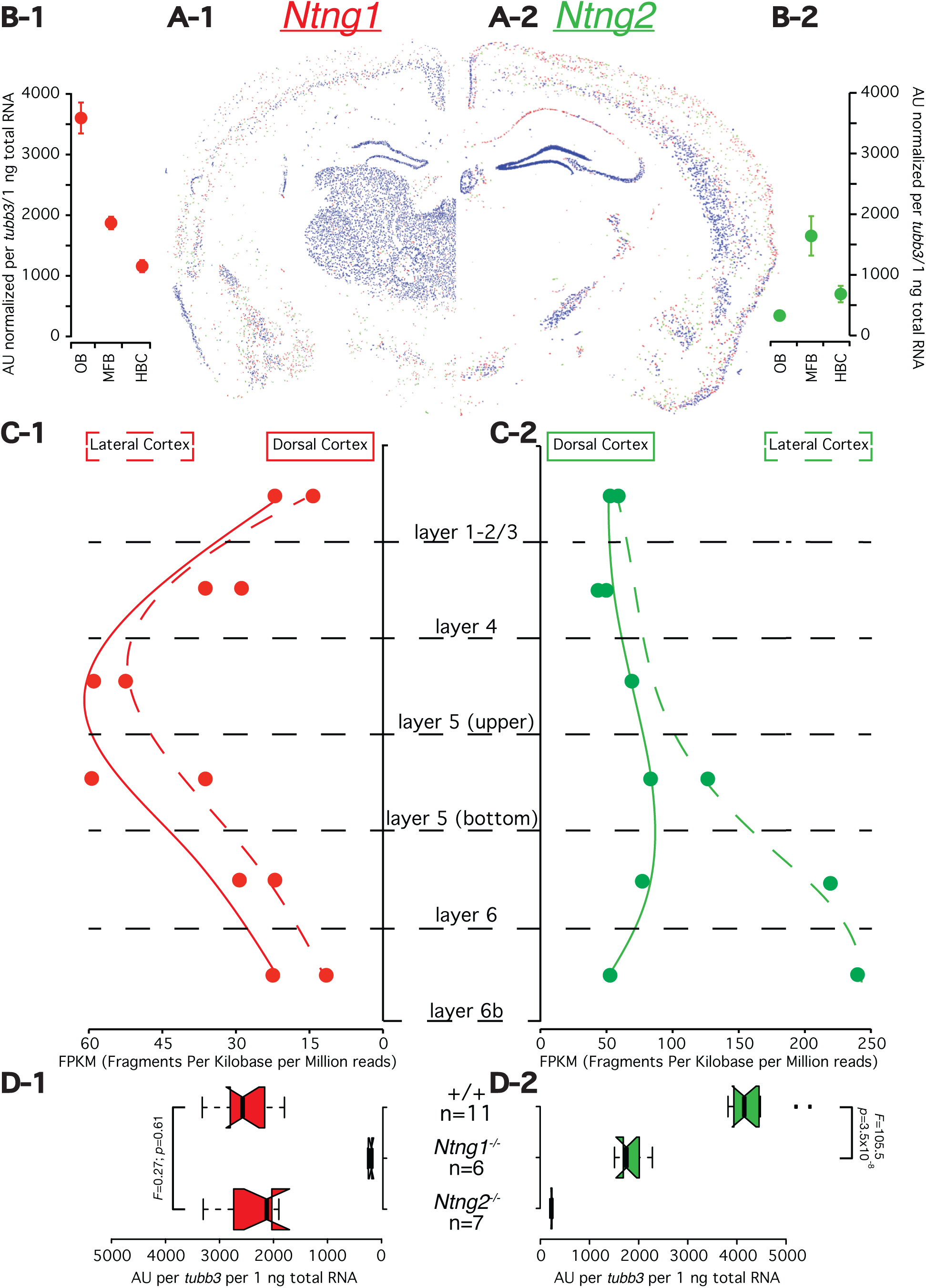
Complementary expression and transcription of Ntng paralogs in the mouse brain. (**A**) In situ hybridization of Ntng1 (left) and Ntng2 (right) of the mouse brain. From Allen Brain Atlas, accession numbers are RP_050607_01_H05 and RP_050810_04_D08, respectively. The expression colors are inverted. (**B**) qRT-PCR of total mRNA for the Ntng paralogs in rough brain fractions of adult naïve male mice (7-8 months old) expressed in the arbitrary units (AU) normalized by *tubb3* per 1 ng of total RNA input per reaction. OB = olfactory bulb; MFB = mid-and front brain; HBC = hindbrain and cerebellum. Data are presented as a mean+SEM, n=6 mice (ST5-1). (**C**) RNA-seq of Ntng mRNA expression within mouse brain cortical layers. The expression model was built based on GSE27243 dataset generated by Belgard *et al*. (51). See Supplementary Methods for the details of data processing, ST5-2, ST5-2_cuff and ST5-2_ireckon (zipped) for the trascriptome assembly. (**D**) Effect of reciprocal genetic background on Ntngparalogs expression level in MFB as detected by qRT-PCR (ST5-3). Senile (20-21 months old) life-long cognitively trained mice have been used (randomly selected from ST1-1 and ST1-2). One-way ANOVA was used for the statistics.

### Genotype prediction based on the phenotype input, the rank

To search for inferences of the genes perturbations on behavioral output we have calculated the probability clustering for each genotype based only on the ranking data input, in genotype-blind manner (fig.6A). The obtained pattern corroborates therelationship between genotypes andassociated with them phenotypes withaffected EF closely resembling the Experimental data (fig.1-3)

### Mouse behavioral phenotypic proximity assessment

To calculate a phenotypic distance between the genotypes comprising a single mixed population we used the obtained ranks and plotted them against the related PVE for each behavioral parameter, generating two linear plots Fig.6B, each representing a single contributing genotype. This let us further to calculate the phenotypic distance (using the classical Euclidean geometry) between the genotypes as the shortest distance between two parallel lines. The obtained geometrical plots are in a full agreement with the experimentally observed behaviors (Figs.1-3) but additionally pinpoint a contribution of each individual parameter sometimes located outside of the main cluster with others, e.g. PreP for the *Ntng1*^-/-^(Fig.6B-1, AI-LCD), OE for the *Ntng1*^-/-^(Fig.6B-2, AI-HCD), and CN for the *Ntng2*^-/-^(Fig.6B-2, WM-LCD). Next, using the Ln rank and its PVE from Fig.4 as (*x,y)* coordinates we have assessed the phenotypic proximity of the *Ntng1*^-/-^and *Ntng2*^-/-^mouse AI and WM phenotypes to the Ln deficit.

**Figure 6.**
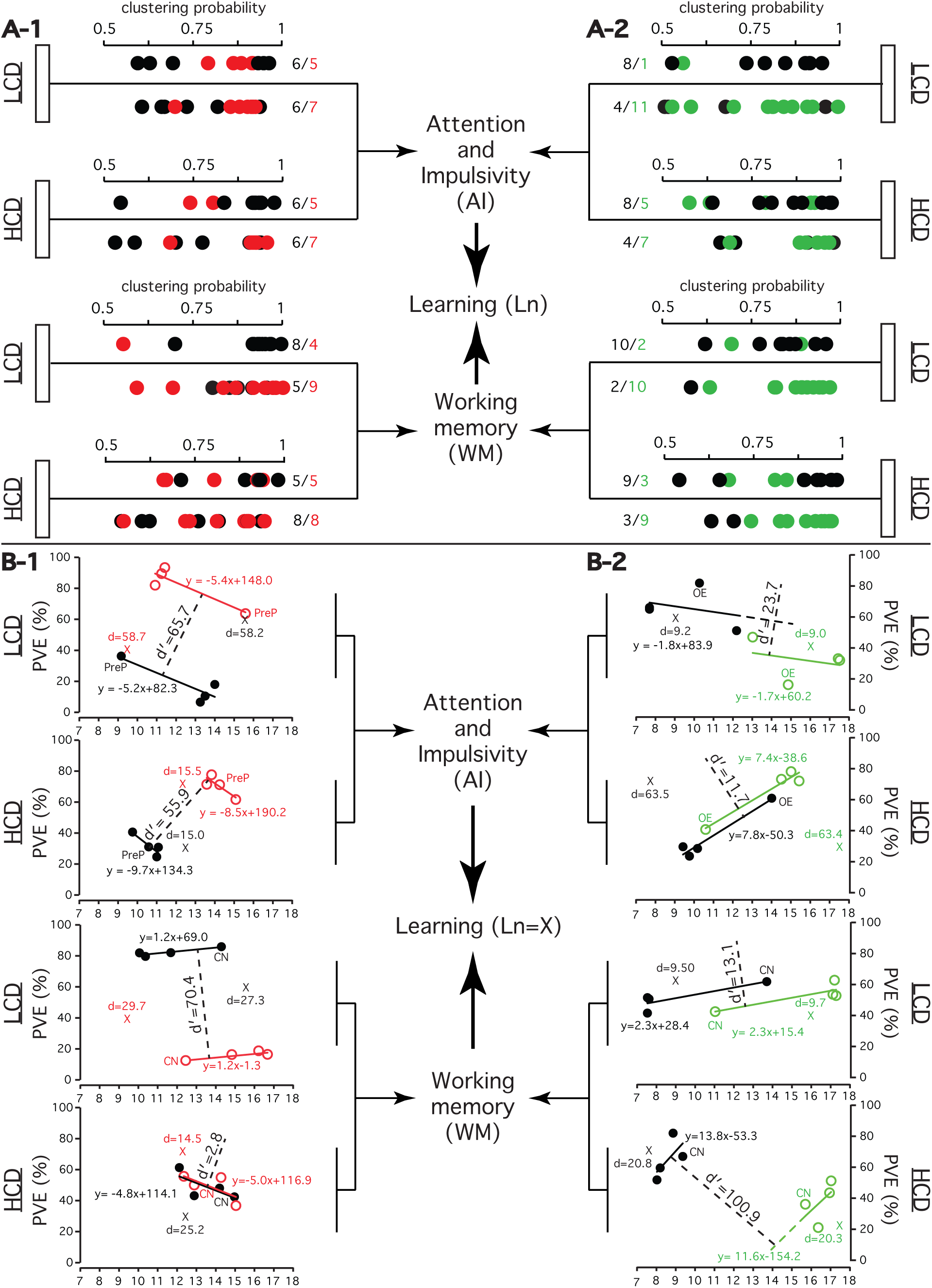
Behavioral phenotypic proximity assessment for the *Ntng1*^-/-^ and *Ntng2*^-/-^ genotypes and their wild littermates by two approaches: Genotype predictions by C-means fuzzy clustering (**A**), and by linear regression plots of genotype-specific rank PVE vs. rank (**B**). **A**. Genotype-phenotype relationship inference. *C*-means fuzzy clustering (Euclidean C-means) was done in a genotype-blind manner as described in SM. See ST6-1 and ST6-2 for the exact values of clustering probabilities. **B**. Descriptive proximity of the operant conditioning learning explained by the AI and WM phenotypes for *Ntng*^-/-^paralogs mice. Distance between the genotypes (d’, dashed line), presented geometrically by the linear equation, was calculated as d’=|c_2_-c_1_|, where ax+by+c=0 (Euclidean geometry). Distance from the learning coordinates (Ln) to the genotype-describing line was calculated as d=|c-(y_Ln_-ax_Ln_)|. For all calculations see ST1-1,1-2 and ST2-1,2-2 “btw genotypes” spreadsheets, summarized in ST7. Rank (x coordinates) and PVE values (y coordinates) are from Fig.1A,E (AI);Fig.2A,E (WM), and Fig.4B (Ln). Data for the *Ntng1*^-/-^/wt population (RAM) are likely to incorporate 7.69% error since they were not normalised to the total number of animals as for the other populations (n=26 for the given case *vs*. n=24 mice for all other three populations).

### Task learning (Ln) ability as an outcome of AI and MW interactions

With the assumption that a shorter distance from the Ln coordinates to the genotype-specific linear plot generates higher likelihood that the given genotype contributes to the Ln associated behavior, we were able to build a relationship graph among the Ln, AI and WM interactions modulated by the cognitive demand (Fig.7A). The dynamics of the *Ntng* gene paralogs hierarchy interaction is presented on Fig.7B, calculated by the reciprocal plug-in of the rank and its PVE for one gene paralog into the linear plot for the other one (ST7).

**Figure 7.**
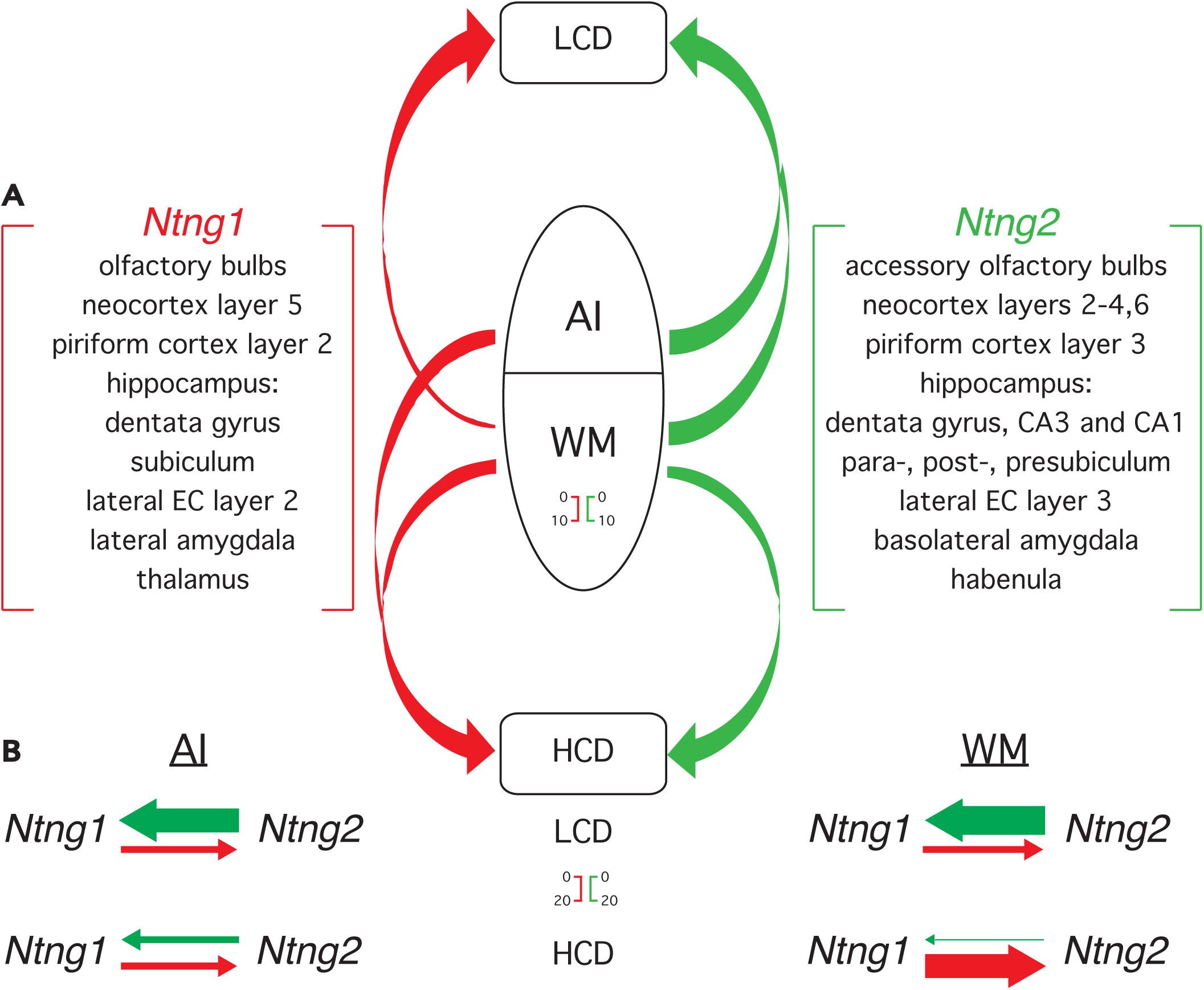
*Ntng* paralogs interaction as a molecular correlate of AI and WM modalities during learning. **A**. AI and WM interactions modulated by the cognitive demand during the operant conditioning learning (Ln), complementary contributed by the differentially expressed Ntng paralogs (see Fig.5 and (52)). **B**. Dynamicity of Ntng paralogs hierarchy interactions under the LCD and HCD contributing to AI and WM. Obtained by reciprocal plug-in of rank and PVE for one Ntng-associated phenotype into the linear equation for another paralog-associated phenotype. Each arrow base width (the scale bar is shown) is expressed in AU and corresponds to (1/d*100) value from Fig.6B (for Ln=X), see ST7. Out of scale arrows (AI-*Ntng1*-LCD and AI-*Ntng2*-HCD) are not shown.

## DISCUSSION

### Inferring genotype-phenotype relationships for the *Ntng* paralogs ablation caused EF perturbations phenotypes

The hierarchy of WM and selective attention interplay has been always a point of fierce debates (21). In the present study we look at this interaction through the prism of mouse operant conditioning learning ability, perturbated by either of *Ntng* gene paralogs ablation (Fig.4). Since it is known that averaging animal behavior across individual subjects (as shown in SF1-1 and SF1-2) may smear out control variables (22), we used rank instead of classical data mean (Figs.1-3) approach and the rank variance (proportion of variance explained, PVE) per genotype, as a measure of difference (23:p.16), to assess the behavioral variability caused by the genetic variations interacting with the experimental demand. To proof any existing inferences between the behavior of *Ntng*^-/-^mice and the ablated gene, we used mouse rank as a randomized dependent variable of the mixed population noting that any non-randomized variables would be only correlational (24). That is, we have presented the mouse behavioral rank distribution as a function of genotype, when one of the *Ntng* paralogs has been genetically inactivated. At the same time, we have tried to elaborate on the statement that the structure of genotype−phenotype map is the matter and not the variance components of the population itself (25). The open question in such genotype-phenotype interaction paradigm is to what degree a genetic variability is capacitive enough to explain the phenotypic variance and the strength of such interaction causality. More specifically, how far the behavioral (whole organism) variability (under the pressure of the growing cognitive demand) represents the neuronal (cellular) variability caused by a gene knockout exerted perturbations.

### Cognitive phenotypes of *Ntng^-/-^*mice

Vertebrate brain-specific presynaptically expressed *Ntng1*^-/-^and *Ntng2*^-/-^ mice do not exhibit gross anatomical or developmental abnormalities (26) rendering them unique models to study brain cognitive functioning in the absence of known “house-keeping” functional distortions, avoiding gene-manipulations-exerted non-causal confounders. Noteworthy the resemblance of *Ntng1*^-/-^and *Ntng2*^-/-^mice behavioral phenotypes with the human schizophrenia subjects behavioral etiology (characterised by the EF control pathologies), both genes have been reportedly associated with (1,2). Two different populations of mice were used for two different behavioral paradigms to avoid the phenomena of learning transfer between the behavioral tests, and, at the same time, to check for the genotype induced phenotypic stability across the different paradigms but sharing the principal underlying component of WM testing. And indeed, slow operant conditioning learning (5-CSRTT) for *Ntng2*^-/-^ mice has been recorded (Fig.4A,B-2) and is explainable by the dysfunction of procedural (working) memory strongly affecting the RAM performance (Fig.2A-H-2).

### Behavior consistency assessment using rank

We have also characterised the behavior of mice as a heterogeneously randomized population through the assessment of rank consistency across the sessions and relative to other parameters (Figs.1-2D,H). Parameters cross-correlation coefficients (r^2, *x* axis) indicate a probability value of how much the rank of a mouse for a certain parameter contributes to the global (total) ranking comprised of all four parameters. If a mouse fails to keep its performance consistent either over the multiple sessions or a parameter, its rank is instantly occupied either by the same or by a littermate of a different genotype, and such event would be dynamically reflected in the r^2. But ranks changes and their permutations may not necessary have any dramatic consequences in the total rank calculations as soon the rank fluctuations are taking place within the same genotype-specific variance boundaries. But they are more reflective of a behavior inconsistency of an individual mouse reflected in the sum of the correlation variances per spatial or session (*y* axis).

### WM deficit driven optimal strategy deprivation for the *Ntng2*^-/-^mice

The global spatial WM deficit for the *Ntng2^-/^*mice has been found robustly expressed across the three RAM parameters (Fig.2AH-2) except for CN (arm choice number during the first 8 arm entries). This parameter represents a strategy development (during LCD) and its optimisation (during HCD) for the maximum reward collection efficiency, akin predictive type behavior of the likelihood of potential success. The fact that the *Ntng2*^-/-^mice outperform their wt littermates in CN (but during the LCD only, Fig.2C-2) reflects the chosen strategy (or a complete lack of any) of a pure random choice of a baited arm to visit, corroborating the global WM deficit (inability for strategic thinking) for the knockouts (evident from the other parameters) but with an opposite valence.

### Paralogs brain expression supporting the behavioral phenotypes

The phenotypic complementarities among the *Ntng1*^-/-^and *Ntng2*^-/-^mice, associated either with the abrogated AI or WM, or both (Fig.3), are supported by the complementary brain expression pattern for these gene paralogs (Fig.5A). If *Ntng1* is expressed mostly in the primary somatosensory gating areas (e.g. OB, thalamus and hypothalamus nuclei, midbrain and medulla, Fig.5A,B-1), *Ntng2* dominates within the cortex (with the skewed expression saturation towards the lateral cortex), hippocampus (HPC), amygdala and claustrum, endopiriform and reticular nuclei, Fig5A-2), pointing the gene role of parsing top-down signals. If the sensory perception, as an entry point into the attentional state, is determined by the strength of the subcortical thalamus-PFC (pre-frontal cortex) pathways (27), the reciprocal interactions between mPFC and HPC are pivotal for the WM functioning (28,29), with the HPC known to encode perceptual representations into memories through the correct attentional states (30). Complementing this,thalamocortical projections are vital for mediating sensation, perception, and consciousness (31-33). It is assumed that WM, despite its distributed nature (34), consists of an executive component spread over the frontal lobes and sensory cortices and interacted by the attention (7,35).

### Brain lamina-specific enrichment and EF control contribution by the *Ntng* paralogs

The emergence of a six-layered neocortex is a known hallmark of the mammalian brain specialization devoted to the EF control (36,37). Both *Ntng* gene paralogs are extensively expressed and mutually sequestered among the separate layers of the cortex (Fig.5C). *Ntng1* is predominantly located in layers 4/5 (Fig.5C-1), probably supporting the arrival of the bottom-up signals (38), while *Ntng2* is located in the superficial layers 2/3 and deeper layers 5/6 (Fig.5C-2), reported as a source of top-down inputs in attention and WM demanding tasks (39). Besides that, *Ntng2* has been also marked as a gene classifier for the granule neurons enriched in the cortex layer 6 (40). In overall, the complementary patterning of the *Ntng* gene paralogs expression supports the laminar-specific distribution of the attention-directed modalities.

### Evidence for the cognitive control taking over the perceptual load

Analysing AI and WM interactions during the task learning (Fig.7A), we have revealed that HCD recruits more *Ntng1* (bottom-up) expressing circuitry comparing to LCD, both by WM and AI, reciprocally replacing the preceding *Ntng2* (top-down) contribution. This potentially points to an augmented peripheral sensory control by upregulating the bottom-up information stream. How to explain such intricacy? Attention exploits a conserved circuitry motif predating the neocortex emergence (41) and WM probably exapts the motor control of forward action modeling also elaborated since ancient times (42). The archaic origin of both modalities limits the fundamental brain resource and constrains information processing, forcing trade-offs among the objects of targeted attention through the top-down control and, possibly, causing a competition between the sensory inputs (43,44) by driving attention at representations in sensory areas where the latter gains entry into WM (7). A model has been proposed that selective attention control is directly linked to the executive control part of the WM system (45) corroborating the statement that attention and WM should no longer be regarded as two separate concepts, see (46) for references. The top-down control of primary sensory processing by higher cortical areas (through the recurrent inputs) has an essential role in sensory perception, as we have just demonstrated. The pervasive penetration of the cognitive control, supported by *Ntng2*, affects the sensory inputs, provided by the *Ntng1* expression.

### An IQ for mice?

The EF control variance attributes to the cognitive performance variance and does not exist independently of general intelligence (47) as a critical determinant of human cognition (48). It is no wonder that, in our hands, the genes affecting WM and attention in mice are the same ones affecting IQ in humans (1) and also associated with a variety of devastating neurological disorders (2) and representing a example of antagonistic functional pleiotropy. The open challenge is to find out to what degree, using *Ntng* gene paralogs as benchmarks, we would be able to conclusively draw on either domain specific or domain general cognitive abilities of mice, or any other non-human animal subjects behavioral intelligence.

Conclusively, *Ntng1* participates in bottom-up, and *Ntng2* in top-down brain information flows support, representing an integrative complementary agreement between perception and cognition as two interacting functions of the brain.

## CONCLUSION

The view of Brain (and Mind) as a modular (domain) system is appealing to evolutionary thinking (49) but is strongly biased towards “the prominence of neural reductionism” (22) pervasively dominating modern neuroscience. There is no strict definition of what a cognitive domain is but it can be viewed as a product of interaction between the top-down and bottom-up underlying neuronal circuits forming bidirectional feedback loops for the executively decisive and sensory information flows, possibly controlling themselves. Genes selectively expressed within such circuits via a non-overlapping pattern represent a tantalizing target to study the cognitive domain makeup and its evolution. An ancient *Ntng* gene duplication (>500 million years ago, preceding the Cambrian explosion) and subsequent co-evolution within the vertebrate genomes made *Ntng* gene paralogs to segregate within the top-down and bottom-up evolving information paths, presumably via subfunctionalisation, under the growing ecological demand (first land/water fish met) but different epistatic environment, both gene paralogs are embedded into. Perception and cognition interplay had eventually culminated in a reflectively subjective representation of the external world, also called consciousness, and explicitly controlled by the EF. Unrevealing molecular correlates of the domain-specific cognitive abilities would help us better understand behavior, e.g. to clearly dissect it on actions (as self-generated thoughts) and responses (cue-induced actions), as a decomposable conjunction supporting the robust functioning of the Brain holistic state.

## MATERIALS AND METHODS

### Animals and behavioral set-ups

Animal rearing and handling experimental procedures were performed in accordance with the guidelines of RIKEN Institutional Animal Care and Experimentation Committee (ethics approval number H292-235(3)). Knockout animals generation and the behavioral set-ups are as described in (18) where the original behavioral datasets have been partially used by us.

### Data analysis

All behavioral and transcriptional data including ranks and PVE calculations with all formulas and graphs are presented in ST1-ST5. The dynamics of the rank change for a specific parameter over the course of study and its congruence with other parameters is depicted on Figs.1-2D,H. No robustness calculations of the rank distribution pattern resistance to a sequential removal of a single behavioral subject were done; neither estimate for the minimal number of the top/bottom ranks representing the obtained pattern; it was empirically decided to be equal to the top and bottom four (Figs.1-2).

### Definition of LCD and HCD

During the 5-CSRTT the cognitive demand was incremented by a shorter cue duration and longer inter-trial intervals, as specified in (18). As for the RAM, the second week of testing (sessions 8-14) was done with half-closed/half-opened doors under the gradually building cognitive demand, internally driven by the behavior optimisation strategy for the maximum likelihood of reward collection, top–down executive–attentional pressure to optimise the behavioral performance outcome, contextually similar to the operant conditioning learning (Ln) of spatial 1 of the 5-CSRTT (Fig.4).

### Real-time qPCR (qRT-PCR)

Primers specifically targeting beginning of each *Ntng* gene paralogs full-length transcripts were designed using Primer3Plus: http://www.bioinformatics.nl/cgibin/primer3plus/primer3plus.cgi. Frozen brains RNA was isolated dissolving the mid-and frontbrain section (MFB) using RNeasy Plus Minikit (Qiagen) and the cDNA was synthesised by the QuantiTect^®^ Reverse Transcription kit (Qiagen) using a mix of the random hexamers and oligodT primers. cDNA synthesised from 1 ng of total RNA was used per a single qRT-PCR reaction. The lack of genomic DNA and the absence of external contaminations were confirmed by the RT-minus reactions. Neuronal-specific *tubb3* transcript (β-tubulinIII) was used as an internal normaliser during the qRT-PCR co-amplifications. The C_t_ values were collected at the threshold value of 0.4 and the arbitrary units (AU) were calculated as:
2^-(Ct (amplicon) - Ct (normalizer))^*10,000

### RNA-seq cortical layers *Ntng* transcriptome reconstruction

The original dataset GSE27243 was generated by Belgard *et al*. (51) and has been reprocessed by Cufflinks and iReckon (ST5-2_cuff.xlsx and ST5-2_ireckon.zip) with the latter being used to build *Ntngs* expression map (ST5-2.xlsx) across the brain cortical layers (Fig.5). See Supplementary Methods for details.

### Fuzzy *C*-Means Clustering

Represents a type of a sequential competitive learning algorithm exhibiting the stochastic approximation problem (50). Was used for the genotype predictions based on the behavioral ranks input under the genotype-blind input conditions. The details are described in the Supplementary Methods.

### Statistics

Correlation coefficients (r^2) were obtained with Excel. One and two-way ANOVA was calculated using StatPlus (AnalystSoft Inc.). Wilcoxon rank sum test was done by Matlab (v.7.9.0 2009b) by the function *ranksum*.

## SUPPLEMENTARY MATERIALS (SM)

Contain Supplementary Methods and References, Supplementary Figures (SF1, SF2), and Supplementary Tables (ST6-7). ST1-ST5 are provided as standalone Excel files, and ST5-2_ireckon is a zip file containing compressed RNA-seq brain layers expression reads.

## ACKNOWLEDGEMENTS

This work was in part supported by the “Funding Program for World-Leading Innovative R&D on Science and Technology (FIRST Program)” initiated by the Council for Science and Technology Policy (CSTP), and KAKENHI 15H04290 from the Japan Society for the Promotion of Science (JSPS).

## SUPPLEMENTARY MATERIALS (SM)

### SUPPLEMENTARY METHODS

RNA-seq data analysis of GSE27243 dataset (Fig. 5C)

FASTQ files containing raw reads were downloaded from [1] originally produced by [2]. The whole dataset consisted of two *groups* (dorsal and lateral cortex areas), and six *samples* (A-F) in each *group*. Each *sample* consisted of 2~3 pairs of FASTQ files -*replicates* taken from one of the six layers of dorsal and lateral part of cortex. Each *replicate* contained approximately 110 million of 50 base pair length paired end reads, where each member of the pair is stored in a separated FASTQ file. Raw reads were filtered by using the set of scripts provided by McDonald Lab [3]. Filtering was necessary for obtaining line-by-line matched set of reads in two files comprising each *replicate*. Filtered FASTQ files were aligned to a mouse reference genome (mm10) by using TopHat v.2.0.8b [4] software which calls for BowTie v.2.1.0 [5], using the following command line options:

*--library-type fr-unstranded --microexon-search --no-coverage-search -G genes.gtf*

Two different approaches for TopHat alignment were used. In the first approach each *replicate* (each pair of FASTQ files) was aligned to the reference genome individually, producing single output bam file per the *replicate*. Second approach employed TopHat’s ability to simultaneously align *replicates* to the same *sample*. This approach yielded a single output bam file per *sample* (12 files in total).

Reference genome sequence, genome indexes and GTF-file containing gene annotations (version mm10 of UCSC build) adapted for use with TopHat/Cufflinks software suite were downloaded from “iGenome” project's web site [6]. TopHat's output was passed then to Cufflinks (v.2.1.1 [ref]) using the next command line:

*$ cufflinks --library-type fr-unstranded -g genes.gtf DCTX_layerA_accepted_hits.bam*

Transcript models generated by the first stage of Cufflinks processing (“transcripts.gtf” files from each *sample*) were combined into one GTF file by using Cuffcompare software – a part of the Cufflinks suite with default options. Resulting file was then used as a reference during the second stage of Cufflinks processing, when it runs in quantification-only mode. Output files generated by Cufflinks (3 files per run, containing FPKM values for genes, isoforms and transcripts expression) were processed with custom Python script. The script combined Cufflinks results from all samples into a single file with the sample labels added and limited the output information to the following genome loci: chr3:109,500,000110,500,000 (for *Ntng1*) and chr2:2,900,0000-2,950,0000 (for *Ntng2*). The resulted file (ST52_cuff.xlsx) was then inspected and analysed manually for the presence of multiple either non-annotated transcripts, or transcripts with varying boundaries or retained after the splicing upstream or downstream introns. However the data presented on Fig.5C represent a sum of all assembled transcripts’ FPKM generated by iReckon [7] allowing for novel (nonannotated) isoforms (Prosselkov et al, unpublished) assembly and quantification (ST52_ireckon.zip), summarized in ST5-2.xlsx. In contrast to Cufflinks, iReckon is able to perform the reads alignments to the specified chromosomes coordinates.

### *C*-Means fuzzy clustering (Fig.6A)

The genotypic distance between phenotypes (Fig.6) was assessed by fuzzy *C*-Means clustering algorithm [8]. For any given number of mice (n=24 or 26, in rows) and their behavioral performances (mouse rank in one of four types of parameters, m=4, in columns) the algorithm calculates cluster number (1 or 2) to which any particular mouse is likely to belong to and probability of the membership. Custom script written in R programming language (package “e1071”, function “cmeans”) was used for the actual calculation. Parameters for the *cmeans* function are as follows:

Number of clusters, or initial values for cluster centers: centers=2

Maximum number of iterations: iter.max=10,000

Distance measure: dist=”euclidean”

Calculation method: method=”cmeans”

Degree of fuzzification: m=2

**Supplementary Figure 1 (SF1).**
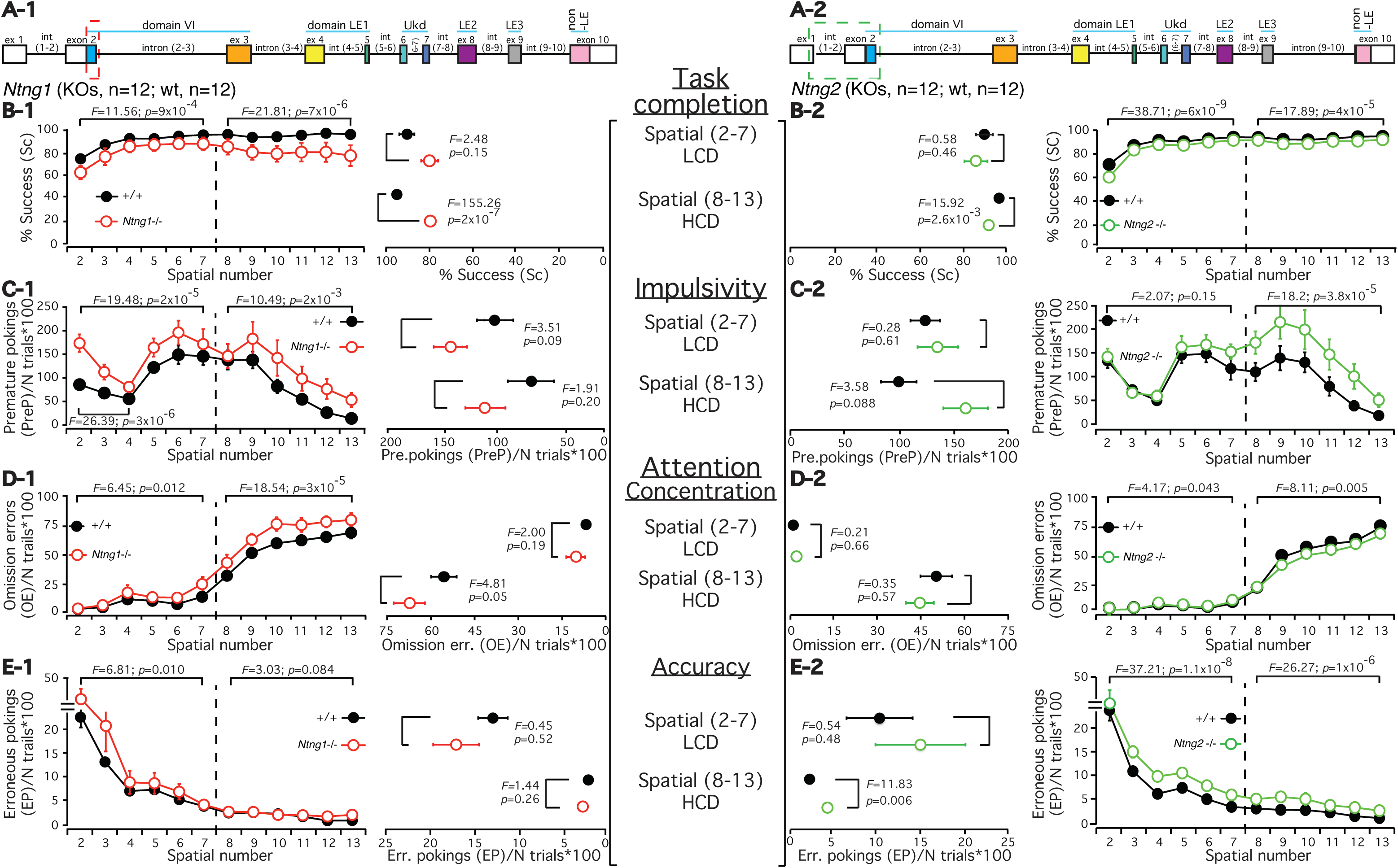
Attention and Impulsivity (AI) estimate for *Ntng1*^-/-^ and *Ntng2*^-/-^ mice by comparison of averaged behavior for 5-CSRTT. A. Exon-intron composition of mouse *Ntng1* and *Ntng2* genes with the removed part for each gene paralog dash-outlined. See (18) for the details of each gene construct design. (**B-E, outmost left and right**) Mouse performance over the spatials (2-13) presented by four behavioral parameters. The dashed line separating spatials (2-7) and (8-13) indicates the data split on the low cognitive demand (LCD) and high cognitive demand (HCD) sessions. (**B-E, middle left and right**) The same data as above but averaged per LCD and HCD sessions. The data for each parameter are presented as a mean±SEM (standarderror of mean) and fully provided in ST1-1 (for *Ntng1*^-/-^) and ST1-2 (for *Ntng2*^-/-^). Two-way and one-way ANOVA were used for statistics.

**Supplementary Figure 2 (SF2).**
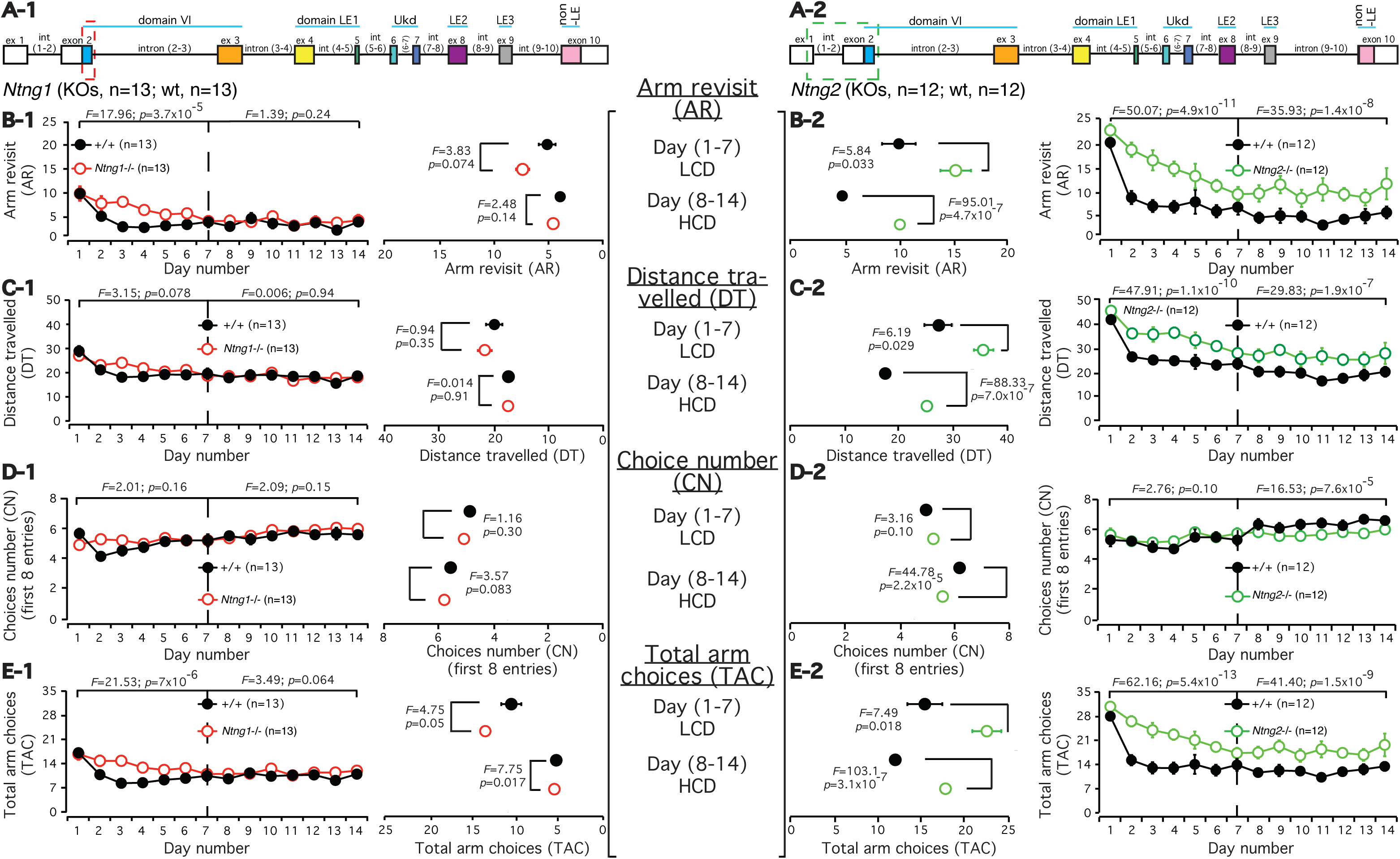
Working memory (WM) estimate for *Ntng1*^-/-^and *Ntng2*^-/-^mice by comparison of averaged behavior for RAM. A. Exon-intron composition of mouse *Ntng1* and *Ntng2* genes with the removed part for each gene paralog dash-outlined. See (18) for the details of each gene construct design. (**B-E, outmost left and right**) Mouse performance over the days (1-14) as presented by four behavioral parameters. The dashed line separating days (1-7) and (8-14) indicates the data split on the low cognitive demand (LCD) and high cognitive demand (HCD) sessions. (**B-E, middle left and right**)The same data as above but averaged per LCD and HCD sessions. The data for each parameter are presented as a mean±SEM (standard error of mean) and fully provided in ST2-1 (for *Ntng1*^-/-)^ and ST2-2 (for *Ntng2*^-/-^). Two-way and one-way ANOVA were used for statistics.

**Supplementary Table 6-1 (ST6-1).**
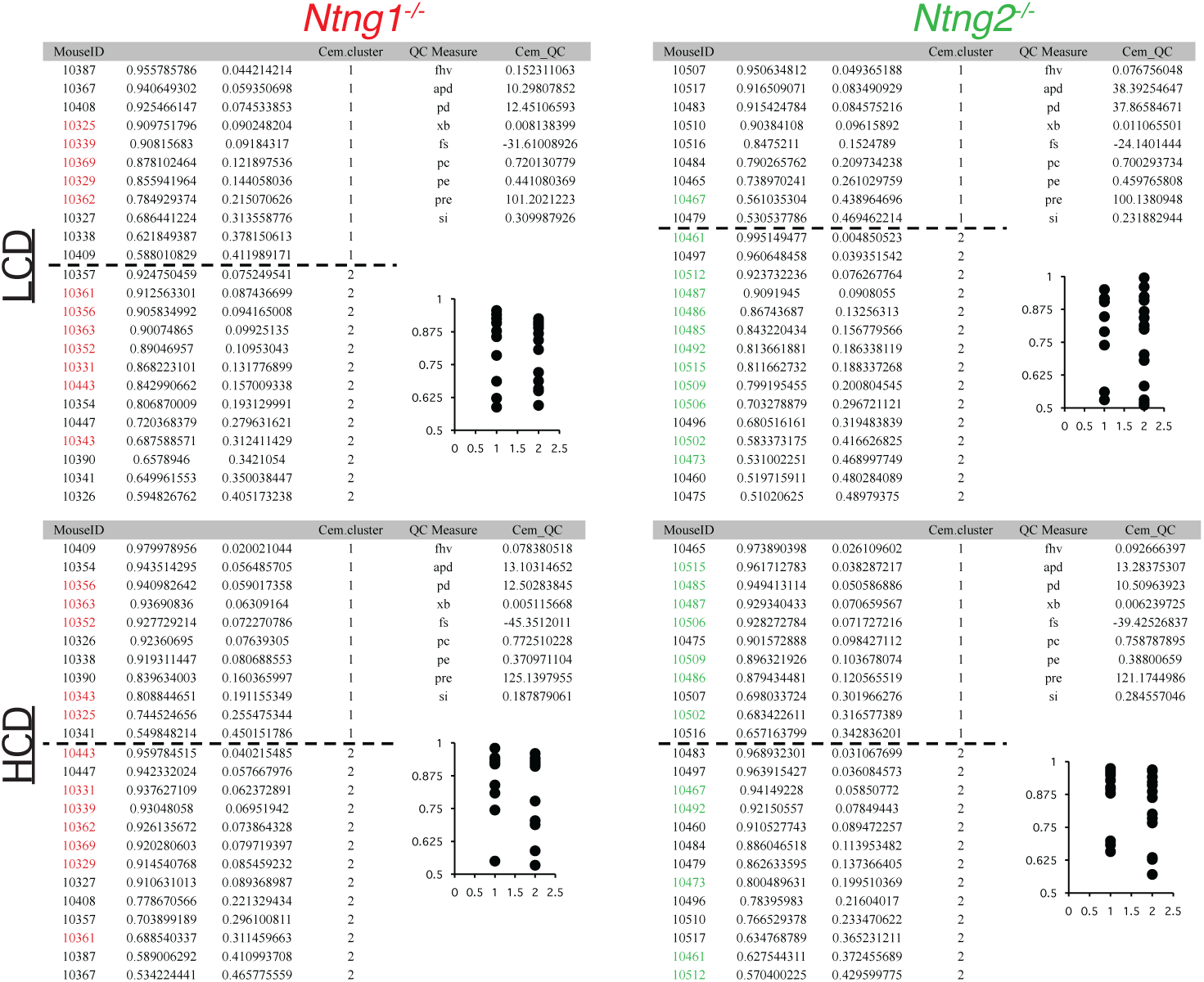
*C*-means fuzzy clustering (Euclidean C-means) rank probabilities for *Ntng1*^-/-^ and *Ntng2*^-/-^ mice performance for the 5-choice serial reaction time task (5-CSRTT).

**Supplementary Table 6-2 (ST6-2).**
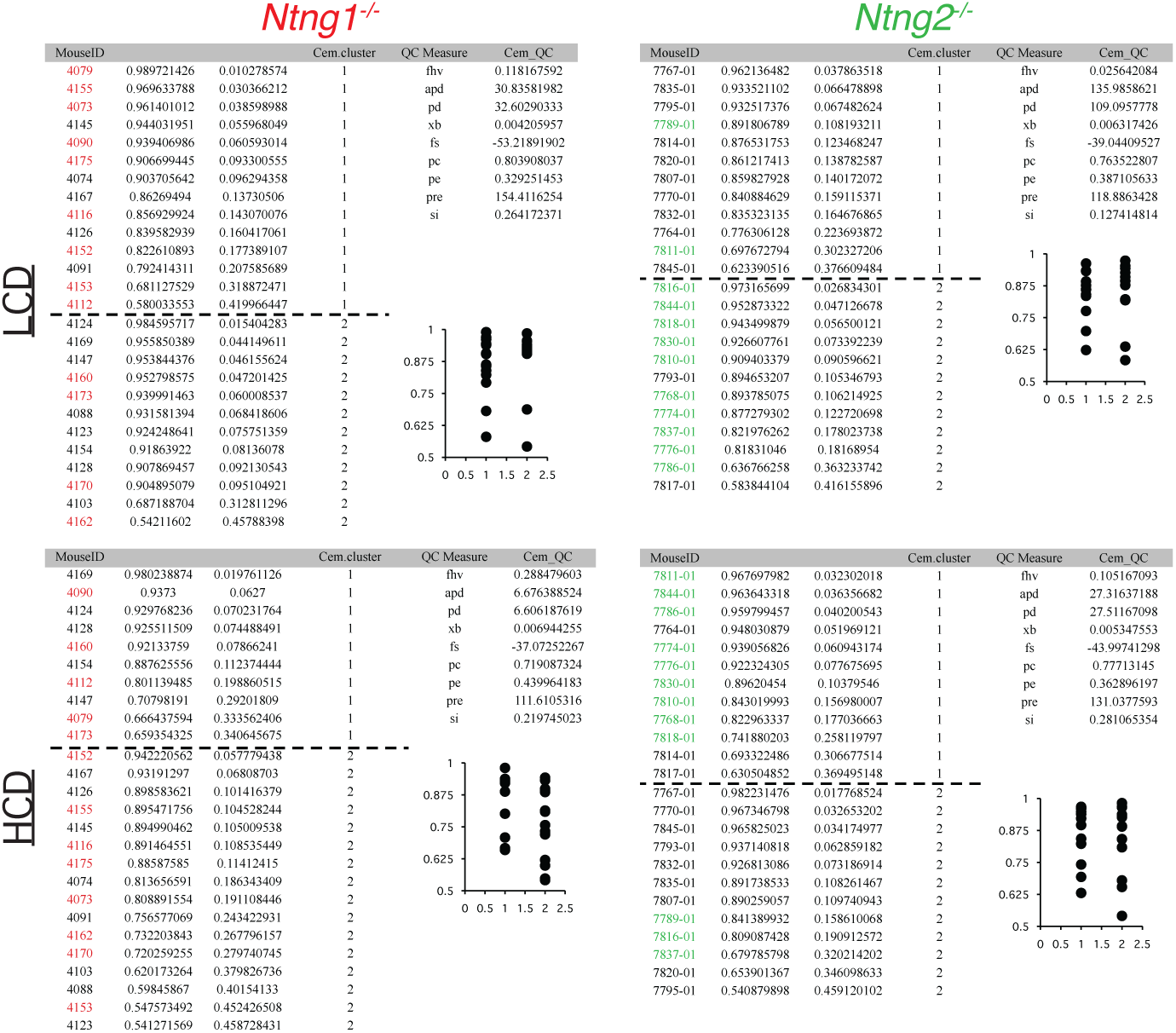
*C*-means fuzzy clustering (Euclidean C-means) rank probabilities for *Ntng1*^-/-^ and *Ntng2*^-/-^ mice performance on the radial arm maze (RAM) task.

**Supplementary Table 7 (ST7).**
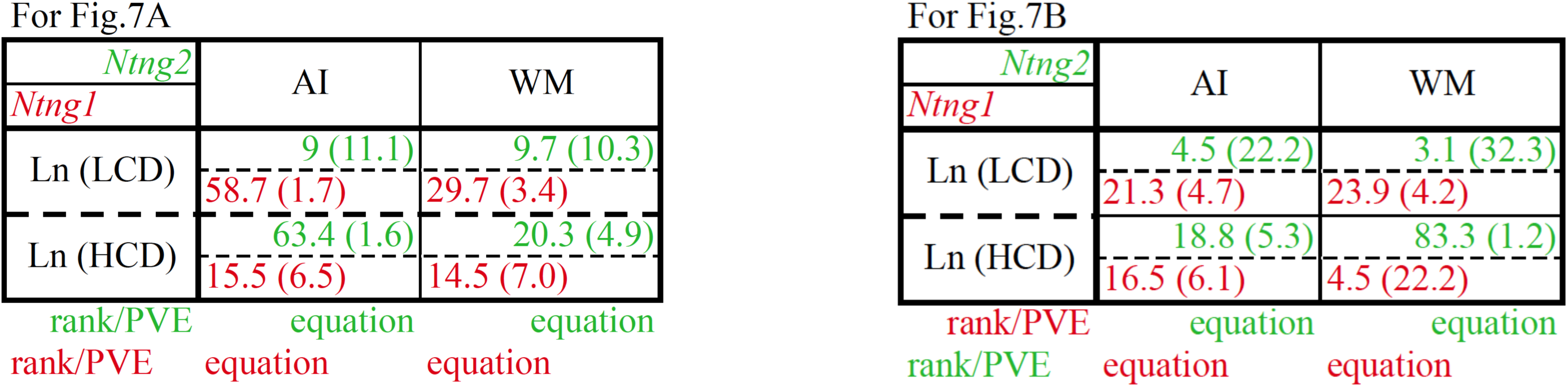
Calculated distance values for the Ntng paralogs, Ln, WM and AI interactions based on the obtained rank and PVE (Figs.1-4)plugged in as linear coordinates (*x*,*y*) into the parameter-derived linear equations from Fig.6B. For Fig.7B the plugging order was reciprocally reversed (shownunder the table). See ST1-1, ST1-2, ST2-1 and ST2-2 (“btw genotypes”) for the distance values calculations and the values describing interactions (in brackets).

